# Immune receptor LAG3 regulates microglia function during Alzheimer’s disease

**DOI:** 10.64898/2026.05.08.723911

**Authors:** Andrew T. Perl, Juan Wu, John D. Dong, Ashley M. Brooks, Andrew R. Yoblinski, Tia T. Vierling, Jian-Liang Li, Dana R. Ruby, Daniel Radzicki, Serena M. Dudek, Jesse D. Cushman, Elizabeta Gjoneska

**Author notes:** **Corresponding Author:** Elizabeta Gjoneska, Ph.D., Neurobiology Laboratory, NIEHS/NIH, Bldg. 101, Rm. F152 (Mail Drop F1-05), 111 T.W. Alexander Drive, Research Triangle Park, NC 27709.

## Abstract

Alzheimer’s Disease (AD) remains the leading cause of dementia globally, yet the exact etiology is not well defined and effective treatments remain unavailable. Here, we report that deletion of the immune checkpoint receptor lymphocyte activation gene 3 (*Lag3*) in a familial AD mouse model, 5xFAD^+^, can rescue molecular, cellular and behavioral phenotypes of neurodegeneration. Specifically, we demonstrate that amyloidosis and microgliosis in the 5xFAD^+^ mice are significantly reduced by *Lag3* deletion. Moreover, we show that *Lag3* deletion attenuates deficits in neurodegeneration-related behavioral phenotypes in the 5xFAD^+^ mice. Transcriptional profiling reveals that *Lag3* deletion suppresses aberrant overexpression of disease associated microglia (DAM) genes in 5xFAD^+^ microglia, effectively restoring homeostatic transcriptional programs. Finally, we observe reduced CD8^+^ T cell infiltration in the brain of 5xFAD^+^ animals after *Lag3* deletion which likely mediates molecular, cellular and behavioral effects resulting from microglia DAM gene activation. Our results highlight a previously unrecognized role for *Lag3* in AD as a critical regulator of microglia function and suggest *Lag3* might be a viable target for novel AD therapeutic interventions.

**Highlights:**

- Immune receptor *Lag3* deletion ameliorates amyloidosis and microgliosis during AD
- *Lag3* deletion attenuates deficits in neurodegeneration-related behavioral phenotypes
- *Lag3* deletion reverses aberrant activation of DAM genes and restores microglia homeostasis
- *Lag3* inhibition presents a viable approach for novel AD therapeutic interventions

## Introduction

Alzheimer’s disease (AD) is the most prevalent neurodegenerative disorder worldwide, characterized by extracellular deposition of amyloid-β (Aβ) plaques, intraneuronal accumulation of hyperphosphorylated tau, and immune cell activation, ultimately leading to synapse loss and neuronal cell death (Selkoe 2001). Because cognitive deficits emerge late in disease progression, elucidating the molecular pathways that impact early stages of pathology remains essential.

Accumulating evidence supports a causal role for the immune system in AD pathogenesis (Hammond *et al*., 2019). For example, aberrant immune activation has been implicated in AD through multiple genome-wide association studies (Jones *et al*., 2010; Jonsson *et al*., 2013), expression quantitative trait loci analyses (Huang *et al*., 2017), RNA sequencing (Keren-Shaul *et al*., 2017; Krasemann *et al*., 2017; Olah *et al*., 2020) and epigenomic profiling (Gjoneska *et al*. 2015). In particular, microglia, the resident immune cell of the central nervous system (CNS), play a central role in regulating immune function during AD progression.

Lymphocyte-activation gene 3 (*Lag3*) encodes an immunoglobulin superfamily transmembrane receptor expressed by a number of immune cells upon activation, including T cells and natural killer cells (Baixeras *et al*., 1992; Lui *et al*., 2018). In the context of peripheral immune cells, *Lag3* has been studied primarily as an immune checkpoint relevant for cancer therapy (Andrews *et al*., 2017). In the CNS, *Lag3* expression is restricted to microglia (Zhang *et al*., 2014; Galatro *et al*., 2017), however, its role in regulating microglia function is not well understood. Studies have suggested a role for LAG3 in Parkinson’s Disease (Freeze *et al*., 2018; Guo *et al*., 2019; Deyell *et al*., 2023), specifically in the binding and cell-to-cell transmission of fibrillar α-synuclein (Mao *et al*., 2016), a protein which, like Aβ, forms amyloid aggregates (Emmenegger *et al.,* 2021). Interestingly, we found that the region just upstream of the *Lag3* promoter is targeted by the transcription factor PU.1, an important regulator of myeloid cell development, which has previously been implicated in AD, suggesting *Lag3* might play a role in mediating PU.1-dependent regulation of microglia function during AD (Gjoneska *et al*., 2015; Huang *et al*., 2017; Ralvenius *et al.,* 2024).

Here, we identify a causal role for *Lag3* in Aβ plaque-driven neurodegeneration. We demonstrate that deletion of *Lag3* decreases Aβ plaque burden, reduces microgliosis and ameliorates behavioral deficits in the 5xFAD^+^ mouse model of AD. Moreover, we show that *Lag3* deletion leads to increased microglial phagocytosis of Aβ plaques and demonstrate the functional effects are likely mediated through *Lag3*-dependent regulation of DAM gene expression. Our findings identify a previously unrecognized function for *Lag3* in maintaining microglia homeostasis and provide insight into the mechanisms underlying the immune system’s contributions to AD, suggesting *Lag3* might be a viable target for novel AD therapeutic interventions.

## Materials and Methods

### Animal use and husbandry

C57BL/6J (B6), *Lag3* knock-out (KO) (B6.129S2-*Lag3*^tm1Doi^/J), and 5xFAD (B6.Cg-Tg(APPSwFlLon,PSEN1*M146L*L286V) 6799Vas/Mmjax) mice were purchased from Jackson Laboratory. B6 and *Lag3* KO mice were bred in-house with 5xFAD mice to produce 5xFAD, wild-type littermate controls, *Lag3* ^tm1Doi^ *^/^* ^tm1Doi^; 5xFAD, and *Lag3* ^tm1Doi^ *^/^* ^tm1Doi^ littermate controls. Genotyping was completed by Transnetyx Inc. (Cordova, TN). Animals were same-sex group-housed and maintained on a 12-hour light/ 12-hour dark cycle with ad libitum access to water and standard chow. For all experiments, both male and female mice were tested and handled separately. All procedures were performed according to NIH guidelines and approved by the National Institute of Environmental Health Sciences’ Animal Care and Use Committee.

### Tissue collection and immunohistochemistry

Mice were deeply anesthetized with sodium pentobarbital (Fatal-Plus, 100 mg/kg) and transcardially perfused with cold 1x phosphate buffered saline (PBS) and 4% paraformaldehyde (PFA) in 1x PBS. Whole brains were removed and fixed overnight at 4°C in 4% PFA in 1x PBS. Following fixation, brains were briefly stored in 1x PBS until sectioning. Using a vibratome, 40 µM thick sections were cut and then stored in 1x PBS containing 0.01% sodium azide for immunohistochemistry analysis.

Following three 1x PBS washes repeated between staining steps, free floating sections were blocked in 10% normal goat serum (NGS), 1% bovine serum albumin (BSA), 0.3 % Triton X-100 in 1x PBS for 1 hour at room temperature. Primary antibodies were diluted in blocking buffer and incubated overnight at 4°C. Next, host-specific Alexa Fluor secondary antibodies (1:1000, Thermo Fisher) were diluted in 0.2 % Triton X-100 in 1x PBS and incubated for 1 hour at room temperature. Sections were stained with the nuclear marker, 4’, 6-diamidino-2-phenylindole (DAPI) for 5 minutes in 1x PBS. Sections were mounted with Fluoromount-G (Southern Biotech) and #1.5 glass coverslips. Primary antibody information is as follows: IBA1 (Cat: HS-234 308, Synaptic Systems, 1:1000), β-Amyloid (D54D2) (Cat: 8243, Cell Signaling Technology), and CD68 (E3O7V) (Cat: 97778, Cell Signaling Technology, 1:1000).

### Image acquisition and analysis

Images were acquired using a Zeiss LSM 780 inverted confocal microscope or Zeiss 880 LSM. For microglia and amyloid-β quantification, maximum intensity z-stacks (*z* = 6 µm, 10x/0.3 NA objective) from n=4 animals per group, one section per animal, were analyzed in FIJI/ImageJ. Microglia number was determined in the anterior hippocampus using ‘Analyze Particles’ command relative to IBA1+ and DAPI+ cells. Amyloid-β was quantified as the percent of area positive for amyloid-β staining within the hippocampus. For amyloid-β engulfment, Z-stacks (*z*= 0.4 µm, 63x/1.4 NA oil objective) from n=5 animals per genotype (except for FAD^+^ n=4 animals), one section per animal, were analyzed using Imaris. Total image volume, IBA1, amyloid-β, and CD68 surfaces were rendered following manual thresholding. Percent engulfment was calculated as previously described (Schafer *et al*., 2014), with normalization to the total volume of IBA and amyloid-β.

### Cell isolation via flow cytometry

Mice were deeply anesthetized with sodium pentobarbital as described above and perfused transcardially with cold 1x PBS. The whole brain was dissected on ice and hippocampi from both hemispheres were isolated and placed in Hibernate A (Gibco) on ice. Tissue was dissociated using an enzymatic neural tissue dissociation kit (Miltenyi Biotec) and mechanical disruption via Pasteur pipettes. Single-cell suspensions were stained for 10 min with anti-CD45-PE (BioLegend, clone 30-F11) and anti-CD11b-APC (BioLegend, clone M1/70). Dead cells were excluded by staining with 7-AAD. Cell sorting was performed on a BD FACS Aria Cell Sorter (BD Biosciences). CD11b^+^ CD45^low^ cells were defined as the microglia population and were sorted directly into RNA lysis buffer (Qiagen) for subsequent RNA sequencing.

For T-cell infiltration, isolated cells were resuspended in FACS buffer and stained for 10 min at 4°C with the following antibodies: anti-CD45-PE (clone 30-F11), anti-CD4-FITC (clone GK1.5), and anti-CD8α-APC (clone 53-6.7), all from BioLegend. Flow cytometric analysis was performed using a BD LSRFortessa instrument (BD Biosciences) and data were analyzed with FlowJo software.

### cDNA library preparation and RNA sequencing

Total RNA from CD11b^+^ CD45^low^ purified hippocampal microglia was extracted using RNeasy Plus Mini Kit (Qiagen) and stored at -80°C. cDNA was generated using the SMART-Seq v4 Ultra Low Input RNA Kit for Sequencing (Takara). The cDNA libraries were prepared using the Nextera XT DNA Library Prep Kit and Nextera XT index Kit (Illumina) and sequenced by the NIEHS Epigenomics and DNA Sequencing Core Facility using a NextSeq 500 NGS system (Illumina).

### RNA-seq analysis

Raw sequencing reads were first assessed using FastQC (version 0.11.7) to evaluate base quality, GC content, and adapter contamination. Reads were then trimmed to remove adaptors and low-quality bases (average quality score < 20) using Trim Galore (version 0.4.4), and aligned to the mouse mm10 reference genome using STAR (version 2.6.0c, Dobin *et al*., 2013) with default parameters. Ambiguous reads that mapped to multiple regions and reads with a MAPQ score less than 10 were removed.

Gene-level quantification was performed using Subread featureCounts (version 2.0.1, Liao *et al.*, 2014) with the mm10 NCBI RefSeq gene annotation (UCSC, v2020-01-10). Genes without read counts in all samples were excluded from further analysis. The remaining genes were analyzed using the edgeR package (version 3.32.1, McCarthy *et al*., 2012) to identify differentially expressed genes between different genotype groups. The Benjamini-Hochberg method was applied to control the false discovery rate. Genes were considered significantly differentially expressed if they met the following criteria: expression detected in at least one sample (RPKM > 1), a fold change over 1.5, and an adjusted p-value < 0.05. Ingenuity Pathway Analysis (IPA; Qiagen) software was used to identify canonical pathways, upstream regulators, and biological functions enriched for differentially expressed genes (DEGs).

Time-series clustering was performed using the Short Time-series Expression Miner (STEM) software (version 1.3.13; Ernst and Bar-Joseph, 2006). Genes were prefiltered to include only those significantly differentially expressed coding genes at the 9-month time point, defined by fold change over 1.5, adjusted p < 0.05, and RPKM > 1 in at least one sample. Fold changes at 3, 6, and 9 months were used as input. Clustering was performed using the STEM algorithm with a maximum of 50 model profiles and 1,000 permutations for significance testing. A minimum correlation of 0.7 was required for gene assignment to model profiles. Statistical significance was assessed by comparing observed gene assignments to random expectation, with profiles considered significant at adjusted p < 0.05. Significant profiles were color-coded (e.g., red, green, blue, yellow) and ranked by p-value. Significant profiles selected for IPA were defined as adjusted p < 0.05 with at least 50 genes.

### Gene set enrichment analysis

Gene set enrichment analysis (GSEA) was performed using the GSEA v4.3.3 desktop application (Subramanian *et al*., 2005). A custom list of 500 DAM-related genes was obtained from (Karen-Shaul *et al*., 2017) and converted to gmt file format for analysis. DeSeq2 v1.44.0 normalized expression values were provided as the expression set in gct file format (Love *et al*., 2014). Gene set labels were permutated 1,000 times and genes were ordered with the Signal2Noise metric.

### Behavior

All animals were handled by the experimenter for three consecutive days, one week before behavioral testing. Animals were acclimated to the room for 30 minutes prior to testing. At the end of each trial, behavioral chambers were cleaned with PREempt disinfecting solution (Contec). A total of 99 mice were run through a behavioral battery consisting of open field, and elevated plus maze, as outlined below, in that order, with each task separated by one week. The experimental design consisted of 10-11 mice per sex, per age, per genotype (WT, 5xFAD^+^, *Lag3*^-/-^, and *Lag3*^-/-^; 5xFAD^+^).

### Open field

Individual mice were placed in an open field box (27 cm x 27 cm) with transparent plastic walls. Automatic recording of spontaneous locomotion began immediately after mice were placed in the center of the testing arena and lasted for 30 minutes. Mice were analyzed for total distance traveled.

### Elevated plus maze

The elevated plus maze consisted of a platform of two 79 cm-long crossed arms raised 47.5 cm above the floor with standard illumination of 25 lux. Animals were acclimated to the room for 60 minutes prior to testing. For each trial, an individual mouse was placed in the cross of the maze and recorded for 5 minutes and their movement was tracked using EthoVision (Noldus). Mice were analyzed for total distance traveled and time spent in the open or closed arms.

### Statistical analysis

All statistical analyses were performed and graphed using Graphpad Prism. For all experiments and data sets, the *p*-value threshold was set to *α* = 0.05. All data are represented as mean ± standard deviation.

Behavioral endpoints were analyzed with multi-factorial ANOVA with Age, Sex, LAG3 status and FAD status as factors. The significance of the main effects and interactions, combined with a priori hypothesis were used to guide subsequent least significant different post-hoc comparisons. This approach helps to limit the number of statistical comparisons made to reduce type I error rate.

### Electrophysiology - Field recordings in acute hippocampal slices

Mice were injected with Fatal Plus (sodium pentobarbital, 50 mg/mL: >100 mg/kg) via intraperitoneal injection, followed by decapitation. The brain was removed and placed in ice-cold, oxygenated cutting solution (in mM: 3.0 KCl, 10.00 D-glucose, 124.00 NaCl, 0.50 CaCl_2_, 7.00 MgCl_2_, 1.25 NaH_2_PO_4_, 26.00 NaHCO_3_) and coronal slices cut at 400 μm on a Leica VT 1200S vibratome. Slices were immediately placed in an interface chamber (Scientific Systems Design Inc, Model#BSC2) set to temperature 32°C, with a working temperature of 31.6°C as monitored by Clampex over the course of the experiment. The chamber was superfused with oxygenated artificial cerebrospinal fluid (aCSF; in mM: 3.0 KCl, 10 D-glucose, 124 NaCl, 2.4 CaCl_2_, 2 MgCl_2_, 1.25 NaH_2_PO_4_, 26 NaHCO_3_, 95% oxygen, 5% CO_2_) at a flow rate of ∼1.4 mL/min. Humidified 95% oxygen, 5% CO_2_ was also supplied from below the interface chamber. Slices were allowed to recover for a minimum of 1 hour before field recordings began.

Pipettes (Sutter Instruments #B150-86-10HP) were pulled using a Sutter Instrument Model P-1000 Flaming/Brown Micropipette Puller resulting in recording electrodes with tip sizes of 1-1.5 µm. Before recording, pipettes were back-filled with oxygenated aCSF. Responses were recorded from an electrode placed in the CA1 stratum radiatum and evoked with a cluster-type stimulating electrode (FHC, CE2C75) also placed in the CA1 stratum radiatum. Data were collected using Clampex 11.1. When a response was found, responses for an input/output (I/O) curve were collected at 25 µA increments until plateau. Subsequent recordings continued at the stimulus amplitude that produced 50% of the maximum evoked synaptic response. Paired pulse recordings were run at 100ms and 500ms intervals for all slices (repeated 3 times, 20 seconds apart).

Data from 2-3 slices were averaged per animal (‘n’) and presented as means ±SEM. The ratio of fiber volley amplitude to synaptic response amplitude was used as an exclusion criterion in that slices with a fiber volley greater than 33% of the response amplitude at the half maximum response were exclude in this data set. The percentage of slices excluded did not differ between genotypes. All recordings were analyzed in Clampfit 10.7 software using the peak amplitude (mV) and initial slope (mV/ms) statistical functions. All experiments on slices were performed by experimenters blinded to mouse genotype through final data analyses.

## Results

### *Lag3* deletion ameliorates microgliosis and amyloidosis in 5xFAD^+^ mice

Amyloid beta accumulation in the neuritic plaques is thought to be a molecular driver of Alzheimer’s disease pathology and progression. Given their close physical association with Aβ plaques, microglia have been implicated in various aspects of plaque pathology, including seeding, compaction, and clearance. To examine the effect of *Lag3* deletion on cellular AD phenotypes, including microgliosis and Aβ plaque accumulation we performed immunofluorescence labeling of Aβ plaques and microglia with the cell-type specific marker Iba1 (Fig. 1A). We observed a 3-fold increase in Iba1-positive microglia in the CA1 area of the hippocampus of 5xFAD^+^ animals at 9 months of age, corresponding to a late stage of disease progression, consistent with the microgliosis phenotype observed during neurodegeneration. Interestingly, microgliosis was significantly ameliorated in 5xFAD^+^ animals lacking *Lag3* (Fig. 1B). Likewise, Aβ plaques accumulation was also observed in the 5xFAD^+^ animals and was most pronounced at 9 months of age (Fig. 1C). Concomitant with the reduction in microgliosis, we also saw a reduction in the β-amyloid plaque burden after *Lag3* deletion (Fig. 1C, Table S1).

**Fig 1.**
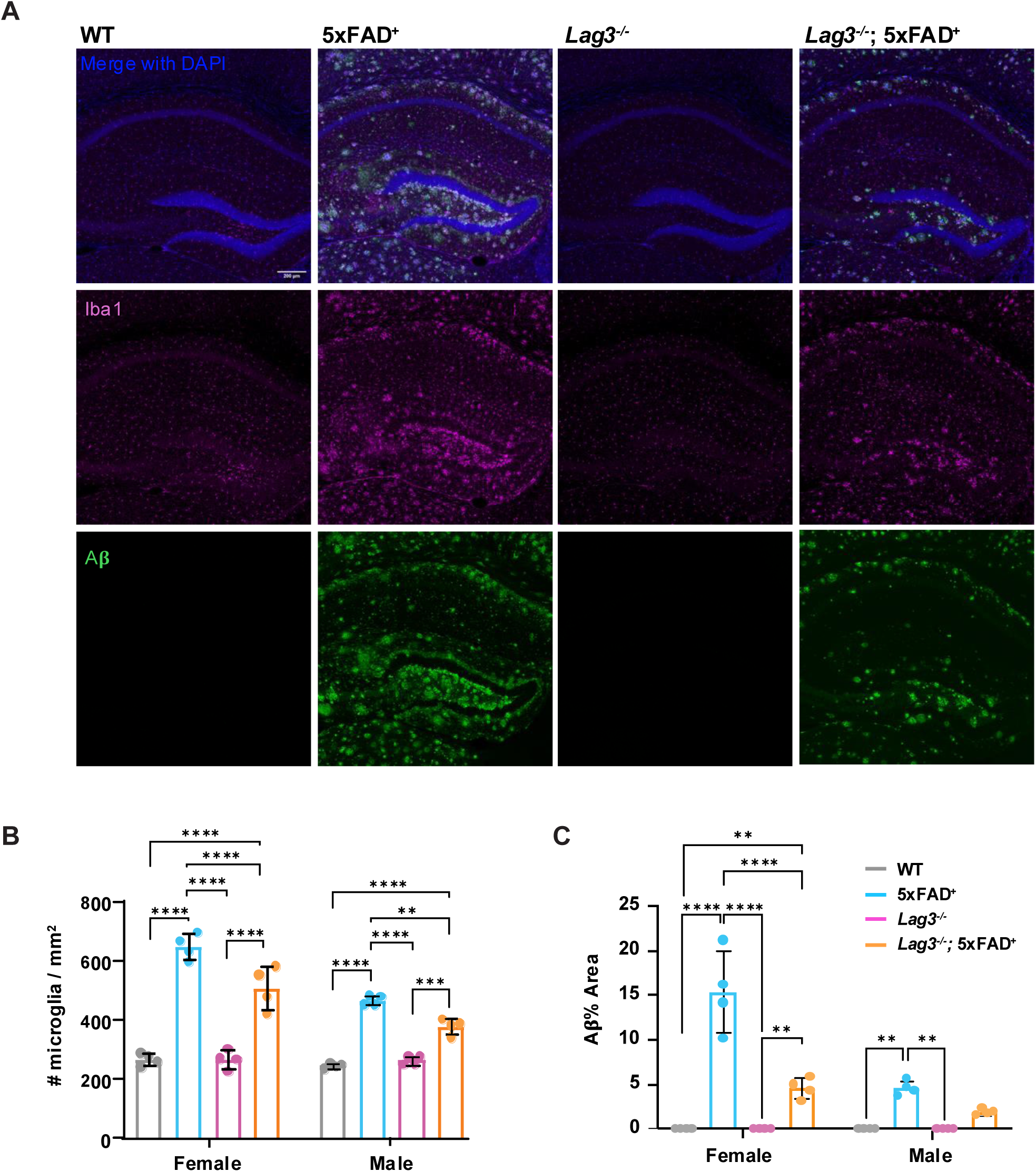
*Lag3* deletion ameliorates microgliosis and amyloidosis in 9-month-old 5xFAD mice. (A), Representative confocal immunofluorescence images of the hippocampus showing Iba1 (magenta) microglia, Aβ (green) plaques, and DAPI (blue). (B), Quantification of microglia cell numbers and (C), percentage of Aβ area in the hippocampus from 9-month-old wild type (WT) (grey), 5xFAD^+^ (cyan), *Lag3^-/-^* (purple), and *Lag3^-/-^*; 5xFAD^+^ (orange) mice (n = 4/group). Data represent mean ± SEM, one-way ANOVA followed by Tukey post-hoc test.

### *Lag3* deletion rescues neurodegeneration-related behavioral deficits in 5xFAD^+^ mice

To determine whether changes in cellular phenotypes translate to behavior, we used open field and elevated plus maze tests (Table S2), based on previously reported neurodegeneration-related phenotypes in these behaviors (Smith and Hopp, 2023; Forner *et al*., 2021). The distance moved in the open field showed an effect of *Lag3* status and FAD status (Fig. 2A, Table S3). Post-hoc analysis revealed increased locomotion in the 5xFAD^+^ mice. This increased locomotion has been observed previously in 5xFAD^+^ mice and is thought to reflect an exploratory habituation deficit consistent with hippocampal impairment (Smith and Hopp, 2023). Interestingly, the exploratory habituation deficits were rescued in the *Lag3^-/-^* mice.

**Fig 2.**
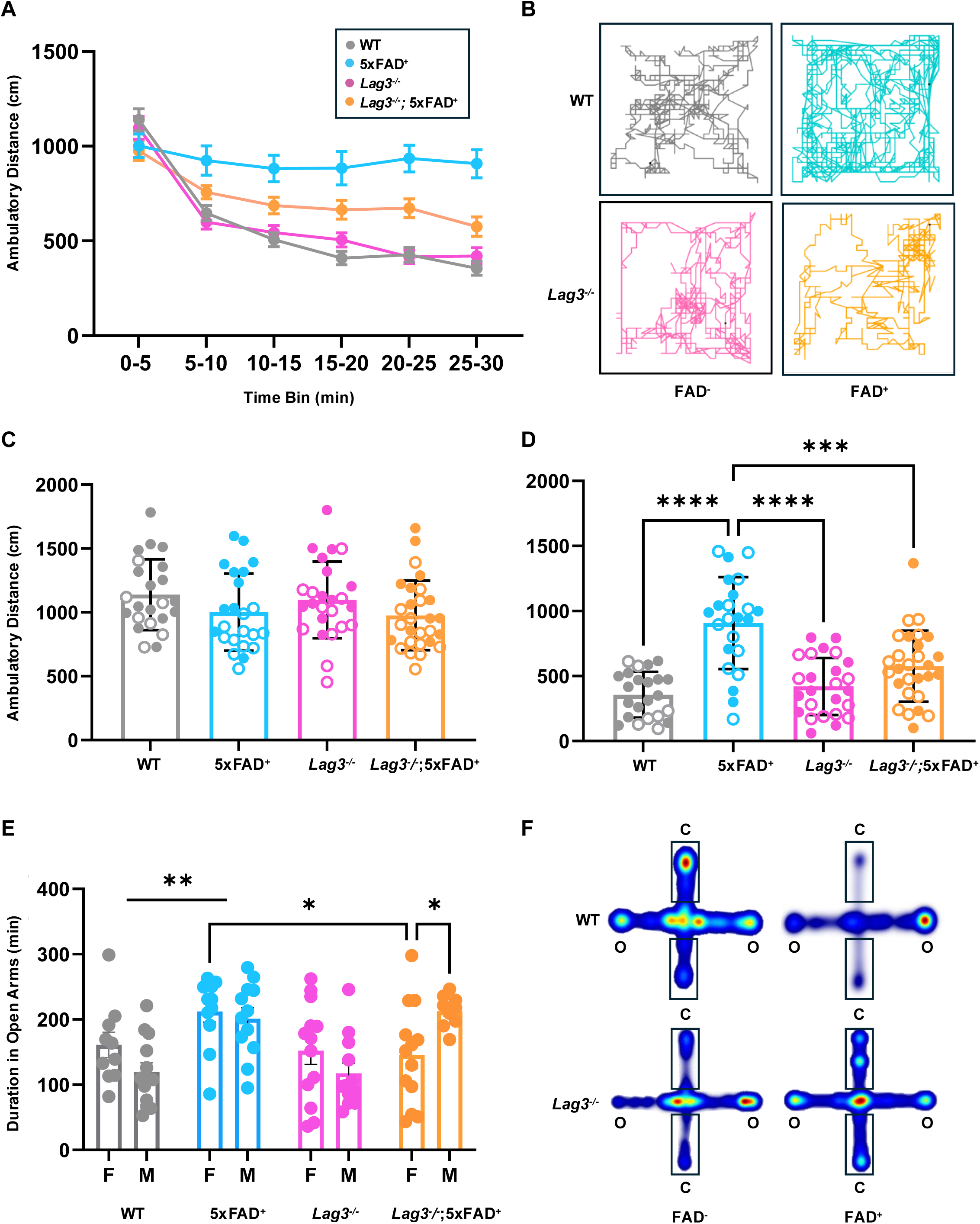
*Lag3* deletion rescues neurodegeneration-related behavioral deficits in 9-month-old WT (grey), 5xFAD^+^ (cyan), *Lag3^-/-^* (purple), and *Lag3^/--^*; 5xFAD^+^ (orange). (A), Quantification of time-dependent ambulatory distance traveled in the open field chamber. (B), Representative races showing movement of animals in an open field chamber over 30 minutes. (C), Distance traveled in the first 5 minutes of the open field showing similar exploratory activity in all of the groups. (D) Distance traveled in the last 5 minutes showing persistent locomotion in the 5xFAD^+^ mice that is rescued by *Lag3* deletion. (E) Quantification of time spent in the open arms of elevated plus maze, indicative of novelty exploratory behavior. (F), Heat maps showing movement of animals in the open (o) vs closed (c) arms of the elevated plus maze. Data represent mean ± SEM. See Supplementary Tables for detailed statistics. Females denoted by open, males by closed circles. Significant difference (p<0.05) denoted by **\***

To determine whether differences in locomotion are related to exploratory behavior, we analyzed the activity pattern in 5-minute bin intervals across the 30-minute open field session. We observed that, while control and *Lag3^-/-^* mice display normal exploratory habituation with reduced locomotion over time, the 5xFAD^+^ mice show persistently elevated locomotion (Fig. 2A, B, C, D). This pattern suggests that 5xFAD^+^ mice fail to exhibit normal context recognition that leads to reduced exploratory drive as the environment becomes more familiar (Wagatsuma *et al*., 2017). This process of familiarization is hypothesized to reflect encoding of environmental attributes via hippocampally-mediated synaptic plasticity (Rudy and O’Reilly, 2001; Krasne *et al*., 2015; Godsil *et al.*, 2005). Impairment of this process is associated with manipulations that disrupt hippocampal function and has been proposed as a hallmark of neurodegeneration-related behavioral deficits in the 5xFAD^+^ model (Smith and Hopp, 2023).

Analysis of the time spent in the open arm of the elevated plus maze revealed effects of FAD status and sex by genotype interaction (Fig. 2E, F, and Table S3). There was an overall elevation of open arm time in the 5xFAD^+^ mice at 9 months, as has been reported previously (Forner *et al*., 2021) and this increase was rescued by *Lag3* deletion specifically in female, but not the male, mice. While the increase in open arm can indicate reduced anxiety, it is likely that it also reflects a similar alteration in novelty-mediated exploration, as we observed in the open field.

These behavioral findings show robust, age-dependent neurodegeneration-related phenotypes in the 5xFAD^+^ mice that are rescued in the *Lag3* deficient mice. The exploratory habituation deficit in the open field is the most robust and sensitive endpoint that is independent of sex. The increased open arm time in the elevated plus maze shows a more complex pattern, with the rescue being specific to females.

Several studies have demonstrated synaptic deficits in the 5xFAD^+^ mice (Forner *et al*., 2021). To determine whether the apparent rescue of pathological changes by *Lag3* knockout is reflected in synaptic function, we recorded field potentials from CA1 stratum radiatum in slices from mice of the four genotypes. Similar to previous studies, we found that the initial slopes and amplitudes (not shown) of synaptic responses from 5xFAD^+^ mice were on average about half of those from WT mice (Supplementary Fig. 1A-E). This was also apparently the case with the *Lag3* deficient 5xFAD^+^. However, we note that any apparent differences between genotype were not statistically significant due to the highly variable responses in our experiments. Interestingly, the *Lag3* knockout by itself seemed to protect from the effects of aging when responses were measured at 9-10 months of age, however again, any possible effect was not significant.

### *Lag3* deletion reverses aberrant gene expression in 5xFAD^+^ mice

To investigate the molecular mechanisms underlying the cellular and behavioral effects of *Lag3* deletion on 5xFAD^+^ phenotypes, we analyzed the transcriptional profiles of hippocampal microglia during various stages of disease progression. At 9 months of age, we found *Lag3* deletion changed expression of 2644 genes relative to WT microglia (absolute value fold change ≥ 1.5; false discovery rate (FDR) < 0.05), with 1409 downregulated and 1235 upregulated genes (Fig. 3A, Table S4). To understand the temporal dynamics of these changes during aging, we compared the differentially expressed genes (DEGs) in *Lag3*^-/-^ versus WT microglia across the 3- , 6-, and 9-month of age and grouped them into 26 distinct clusters using a K-means clustering algorithm (Supplementary Fig. 2, Table S5). Notably, DEGs in 2 clusters (profiles of 12 and 17) showed progressive downregulation, whereas 3 clusters (profiles of 8, 13, and 16) exhibited gradual upregulation over time. Pathway analysis of these clusters revealed that *Lag3* deletion enriched pathways are associated with phagocytosis, mitochondrial-related processes, and cell cycle regulation (Fig. 3B).

**Fig 3.**
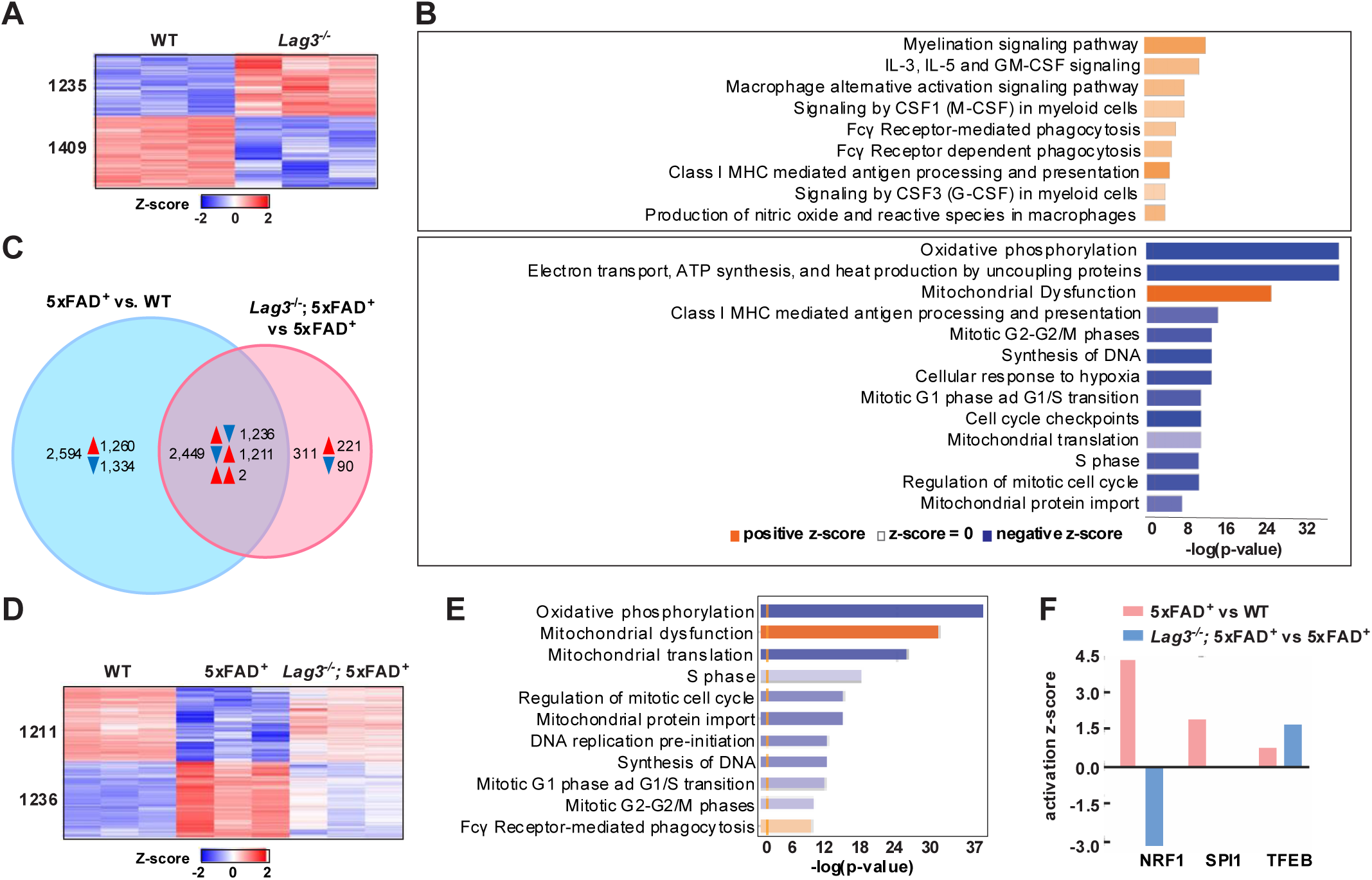
*Lag3* deletion reverses aberrant expression of genes important for mitochondrial, cell cycle, and receptor-mediated phagocytosis in 9-month-old 5xFAD microglia. (A), Heatmap of differentially expressed genes (DEGs) from *Lag3^-/-^* vs WT microglia, with scale representing z-score values. (B), Select canonical pathways altered in *Lag3*^-/-^ vs WT microglia. (C), Venn Diagram showing DEGs in hippocampal microglia between 5xFAD^+^ vs WT and *Lag3*^-/-^; 5xFAD^+^ vs 5xFAD^+^ mice (n = 3/group). (D), Heatmap of overlapping DEGs showing reversed aberrant expression between 5xFAD^+^ vs WT and *Lag3*^-/-^; 5xFAD^+^ vs 5xFAD^+^ mice (n = 3/group), with scale representing z-score values. (E), Selected canonical pathways altered in *Lag3^-/-^*; 5xFAD^+^ vs 5xFAD^+^ microglia. (F), Bar graph depicting selected transcription factors involved in mitochondrial-related processes and immune responses: NRF1 (oxidative phosphorylation), PU.1 (myeloid cell differentiation and immune regulation), and TFEB (phagocytosis regulation).

Comparisons of 5xFAD^+^ versus WT and *Lag3*^-/-^; 5xFAD^+^ versus 5xFAD^+^ hippocampal microglia revealed that *Lag3* deletion reversed aberrant expression of 2447 DEGs observed in 5xFAD^+^ microglia, with 1236 downregulated and 1211 upregulated genes (Fig. 3C, D and Table S4). Canonical pathway analysis further demonstrated that *Lag3* deletion reinstated the expression of genes critical for phagocytosis, mitochondrial function, and cell cycle regulation (Figure 3E). Transcription factor enrichment revealed the involvement of NRF1 (regulating oxidative phosphorylation), PU.1 (myeloid linage cell differentiation), and TFEB (phagocytosis) in *Lag3* deficient 5xFAD^+^ mice (Fig. 3F). To examine the functional impact of *Lag3* deletion on microglia phagocytosis we stained Iba1^+^ microglia for Aβ_42_ as well as the lysosomal protein CD68, a marker of phagocytosis. Three-dimensional reconstruction of Aβ_42_ and CD68 immunofluorescence using Imaris demonstrated that *Lag3* deletion reduced the ratio of Aβ_42_^+^ in both Iba1-positive microglia (Fig. 4A, B) and CD68^+^ phagosomes (Fig. 4A, C), suggesting that reduced plaque burden in the *Lag3* deficient 5xFAD^+^ mice is a result of more efficient Aβ processing by *Lag3^-/-^* microglia. These findings underscore a critical role for *Lag3* in the regulation of phagocytosis in microglia, which may contribute to ameliorated neurodegenerative phenotypes and improved cognitive performance in the 5xFAD^+^ mice.

**Fig 4.**
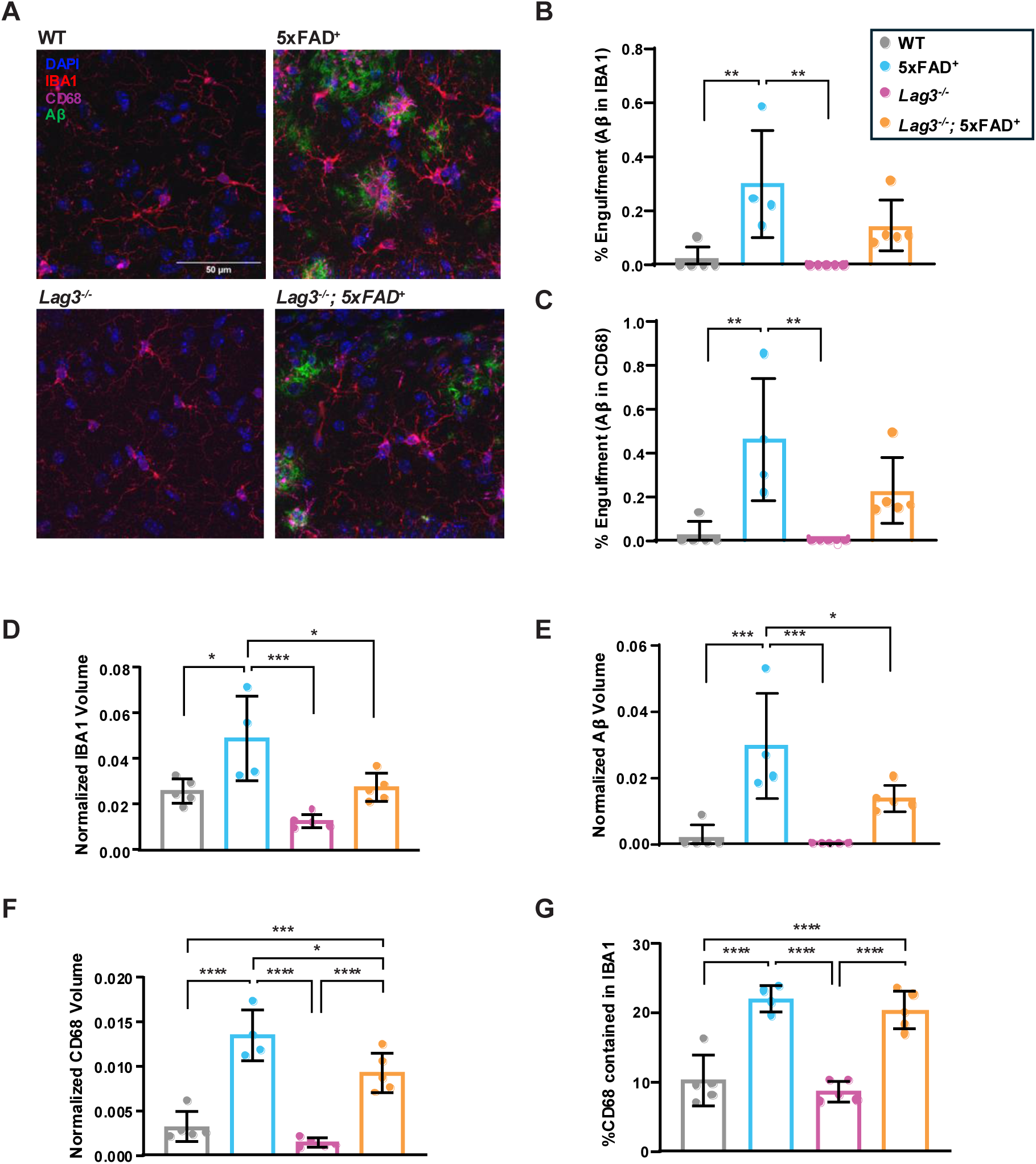
*Lag3* deletion improves Aβ processing in 9-month-old 5xFAD mice. (A) Confocal images showing Aβ (green) internalization by Iba1 (red), and CD68 (purple) positive cells (A), and corresponding quantification of % Aβ engulfment in (B) Iba1 positive microglia and (C) CD68 positive microglia for WT, 5xFAD^+^, *Lag3^-/-^*, and *Lag3^-/-^;* 5xFAD^+^ hippocampus (n = 5 mice/group, except for 5xFAD^+^ n = 4 mice/group). Corresponding normalization for (D) Iba1 volume, (E) Aβ volume, (F) CD68 volume and (G) %CD68 contained in Iba1 positive cells. Data represent mean ± SEM, one-way ANOVA followed by Tukey post-hoc test.

### *Lag3* deletion reduces CD8+ T cell infiltration and suppresses activation of disease-associated microglia signature genes in the brains of 5xFAD^+^ mice

Neurodegenerative microglia phenotype (MGnD) also referred to as disease-associated microglia (DAM) has been previously characterized as a subset of microglia exhibiting gene expression signatures associated with the conversion of homeostatic microglia to disease state during neurodegeneration (Karen-Shaul *et al*., 2017; Krasemann *et al*., 2017). To examine the enrichment of DAM-signature genes in the *Lag3*^-/-^ animals, we performed a gene-set enrichment analysis (GSEA) using the DAM gene set (Table S6). We observed enrichment of DAM-signature genes among the upregulated genes in the 5xFAD^+^ relative to the WT animals at 9 months of age. The same gene signatures were also upregulated in the 5xFAD^+^ animals relative to the *Lag3* deficient 5xFAD^+^ counterparts, revealing that 5xFAD^+^ DAM gene signatures after *Lag3* deletion resemble those of WT animals (Fig. 5A, B). The correlation of rank metric scores between 5xFAD^+^ vs. *Lag3^-/-^*; 5xFAD^+^ and 5xFAD^+^ vs. WT was positive (R = 0.73, p-value = 1.111682e-58), indicating a similar DAM gene ranking in each comparison (Fig. 5C). These results reveal that the loss of microglia homeostatic function observed during neurodegeneration in the 5xFAD^+^ animals is reversed after *Lag3* deletion.

**Fig 5.**
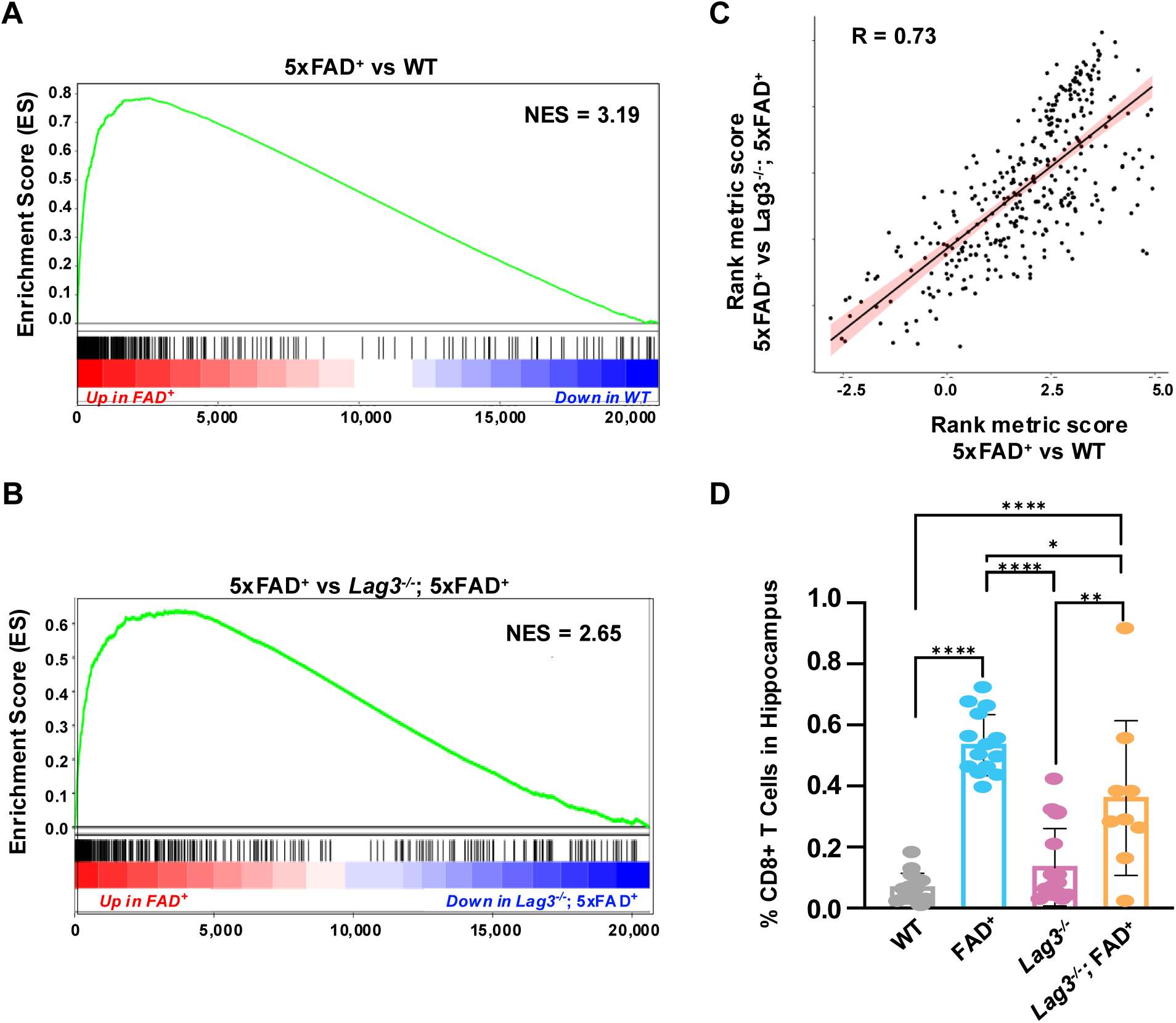
*Lag3* deletion suppresses microglia activation and CD8+ T cell infiltration in the brains of 9-month-old 5xFAD^+^ mice. Gene Set Enrichment Analysis plots of Disease Associated Microglia (DAM) genes display running Enrichment score (ES) and z-score transformed expression values arranged according to ranked gene order for (A), FAD^+^ vs. WT and (B), FAD^+^ vs. *Lag3^-/-^;* 5xFAD^+^ comparisons. (C), Scatterplot comparing GSEA rank metric scores for DAM genes between FAD^+^ vs. WT and FAD^+^ vs. *Lag3^-/-^;* 5xFAD^+^ comparisons indicating high correlation (correlation ratio of R = 0.73, p-value = 1.111682e-58). (D), Quantification of FACS sorted CD8+ positive T cells from the hippocampi of WT (grey), 5xFAD^+^ (cyan), *Lag3^-/-^* (purple), and *Lag3^-/-^*; 5xFAD^+^ (orange) mice. Data represent mean ± SEM, two-way ANOVA followed by Tukey post-hoc test.

To uncover the mechanism mediating the conversion of homeostatic microglia to DAM in the 5xFAD^+^ animals, we measured the number of infiltrating peripheral immune CD8^+^ T-cells in the hippocampus by FACs. We observed a small, but significant increase (0.5%) of CD8^+^ T-cells in the 5xFAD^+^ animals relative to WT controls at 9 months of age and this increase was ameliorated by *Lag3* deletion in the 5xFAD^+^ animals (Fig. 5D, Table S7). This suggests that the microglia conversion to a disease phenotype might be mediated through *Lag3*-facilitated CD8^+^ T-cell infiltration.

## Discussion

In this study, we demonstrate that deletion of the *Lag3* immune checkpoint receptor in the 5xFAD^+^ mouse model of AD rescues both cellular and behavioral phenotypes associated with neurodegeneration and reprograms microglia to a neuroprotective phenotype characterized by reduced microgliosis. These reprogrammed microglia contribute to reduction in Aβ plaque burden and improved cognitive performance in 5×FAD^+^ mice. Transcriptional profiling indicates that the protective effects of *Lag3* deletion are mediated via reversal of aberrant gene expression that typically occurs during AD progression. Specifically, *Lag3* deletion suppresses activation of DAM signature genes seen during AD and restores homeostatic transcriptional programs, thereby preserving the neuroprotective role of microglia in amyloid-β pathology.

While the function of additional immune-checkpoint molecules such as PD1, PD-L1 and TIM3, has previously been investigated in mouse models of AD (Rosenzweig *et al*., 2019; Baruch *et al.*, 2016; Kimura *et al*., 2025), this is the first study to implicate the LAG3 immune checkpoint receptor in AD pathology, thereby establishing it as a viable target for novel AD therapeutic interventions. Systemic blockade of PD1–PD-L1 signaling has been shown to attenuate both, Aβ and tau pathologies by increasing the recruitment of monocyte-derived macrophages and enhancing the frequency of CD4^+^ regulatory T cell in the brain (Chen *et al*., 2023). In aged mice, PD1 blockade additionally mitigates the accumulation of senescent cells by promoting CD8^+^ T cell activation. These CD8^+^ T cells exert a protective role against Aβ pathology through T cell–microglia interaction-dependent mechanisms (Su *et al*., 2023). Conversely, reduced microglia-mediated T cell infiltration has been shown to ameliorate brain atrophy and tau pathology (Chen *et al.*, 2023). Likewise, deletion of TIM-3 exerts neuroprotective effects by uncoupling pro-phagocytic from pro-inflammatory transcriptional programs in microglia, thereby promoting a protective microglia state (Kimura *et al*., 2025). Collectively, these findings highlight the multifaceted roles of immune checkpoint proteins in neurodegeneration. Here, we demonstrate that LAG3 checkpoint inhibition alleviates Aβ pathology by modifying transcriptional programs and suppressing aberrant expression of DAM genes. This effect is likely mediated through the diminished infiltration of CD8^+^ T cells observed in the absence LAG3 in the brains of 5xFAD^+^ animals, suggesting that LAG3 signaling pathways are important for regulation of T cell recruitment via peripheral immune cell-microglia interactions in AD. Interestingly, much like LAG3, deletion of the TIM-3 checkpoint inhibitor has been shown to exert neuroprotective effects by uncoupling pro-phagocytic from pro-inflammatory transcriptional programs in microglia, thereby promoting a protective microglia state (Kimura *et al*., 2025). Collectively, these findings highlight the multifaceted roles of immune checkpoint proteins in neurodegeneration.

Previously published temporal single-cell RNA-seq analysis of hippocampal microglia in a mouse model of AD demonstrated that MHCII genes (*H2-Ab1* and *H2-Aa*), which encode LAG3 checkpoint receptor ligands, are also upregulated during late stages of AD progression (Mathys *et al*., 2017) . Similar increases in MHCII gene expression, as well as upregulation of additional LAG3 ligands, such as GAL-3, has been reported in human postmortem brains of individuals with AD (Boza-Serrano *et al*., 2022). Therefore, it is likely that T cell-microglia interactions in the brain, mediated through LAG*3* ligands such as MHCII and GAL-3, may be important for T-cell recruitment and T cell-mediated microglia activation during AD. As such, it would be of particular interest to investigate whether deletion of LAG3 ligands can similarly exert neuroprotective effects and attenuate AD pathology. Collectively, our results highlight an important role of *Lag3* in regulating microglial function and support its inhibition as a targeted strategy to enhance plaque clearance and alleviate AD pathology.

## CRediT authorship contribution statement

EG, ATP and ARY conceived of the study. ATP, JDD, ARY, JW and TTV performed experiments. JL, AMB, and JW, analyzed sequencing data. ATP, JW, HRR and EG wrote the manuscript with input from JDC, and SMD.

## Supporting information

Table S1. Compiled IHC Data and Stats

Table S2. Compiled Behavior Data

Table S3. Open Field Stats

Table S4. 9m F RNAseq DEGs

Table S5. STEM

Table S6. DAM GSEA Rank Metrics

Table S7. Compiled T cell Data and Stats

## Acknowledgments

The authors would like to thank Lisa Angermeier for colony management, and Carl Bortner and Maria Sifre for help with flow cytometry, the staff the NIEHS Fluorescence Microscopy and Imaging Center for their help with imaging and the staff at the Epigenomics and DNA Sequencing Core for help with sequencing.

## Data availability

Raw RNA-sequencing data are available from the NCBI Gene Expression Omnibus (GEO) database under accession number GSE307127.

