## Supplementary Figures for "Immune receptor LAG3 regulates microglia function during Alzheimer’s disease"

**Fig S1. Synaptic properties in slices from mice of WT, 5xFAD, *Lag3*$^{-/-}$, and *Lag3*$^{-/-}$; 5xFAD mice.** Data are shown from 6-month (left) or 9.5-month old (right) mice. (A-E) No significant differences in synaptic effectiveness were observed, as measured by input/output (I/O) curves. Plotted are initial slopes (μA/msec) of the field excitatory postsynaptic potentials (fEPSPs) recorded in the CA1 stratum radiatum of WT, 5xFAD, *Lag3*$^{-/-}$, and *Lag3*$^{-/-}$; 5xFAD mice at 6 months (A) and 9.5 months (B) of age. Slopes at 250 μA are compared between groups at 6 months (C) and 9.5 months (D), and additionally, slopes at 400 μA are compared at 9.5 months (E). Insets are individual traces from *Lag3*$^{-/-}$; 5xFAD$^{+}$ mice. Although responses from 5xFAD$^{+}$ mice (cyan) showed a strong trend toward being reduced compared to those of WT (grey and pink), the effect showed no evidence of rescue by *Lag3*$^{-/-}$; (*Lag3*$^{-/-}$; 5xFAD$^{+}$, orange) at 6 months (C). Note that there was an effect of genotype at 250 μA in (C) using a Kruskal-Wallis test (p = .0362), but multiple comparisons tests showed no significant differences between any two groups. No significant difference between groups was observed at 9.5 months when comparing responses to either a 250 μA stimulation (D) [F(3,23)=1.489, p =.2438] or a 400 μA stimulation (E) [F=(3, 23) = 1.805, p = .1743], one-way ANOVA test. No significant difference between groups was observed at 9.5 months when comparing responses to either a 250 μA stimulation (D) [F(3,23)=1.489, p =.2438] or a 400 μA stimulation (E) [F=(3, 23) = 1.805, p = .1743], one-way ANOVA test.

**Fig S2. Time-series gene expression clusters reveal temporal dynamics of microglial gene expression changes during AD progression.** Time-series analysis of gene set expression profiles in *Lag3*$^{-/-}$ versus WT microglia at 3-, 6-, and 9-months of age. Each box represents a distinct temporal expression pattern (profile), with the profile ID shown in the top left corner and the associated enrichment p-value at the bottom. Colored boxes indicate statistically significant profiles (adjusted $p < 0.05$): red for downregulated, green for upregulated. Profiles are ordered by number of genes. Two significant profiles (clusters 17 and 12) contained genes that were progressively down-regulated, while three significant profiles (clusters 8, 13, and 16) comprised genes that were up-regulated at 9-months relative to earlier time points.

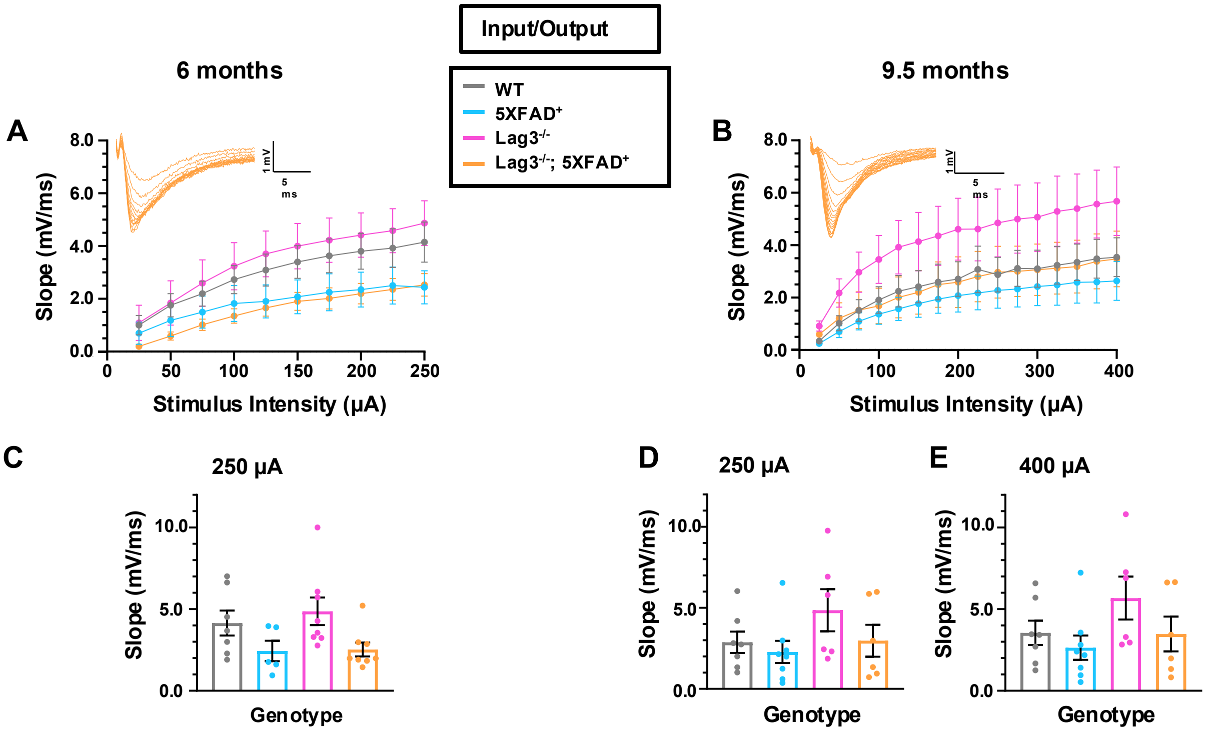
Input/Output
WT
5XFAD+
Lag3-/-
Lag3-/-; 5XFAD+
6 months
9.5 months
A
B
Slope (mV/ms)
Stimulus Intensity (µA)
1 mV
5 ms
C
250 µA
D
250 µA
E
400 µA
Genotype

Figure S1

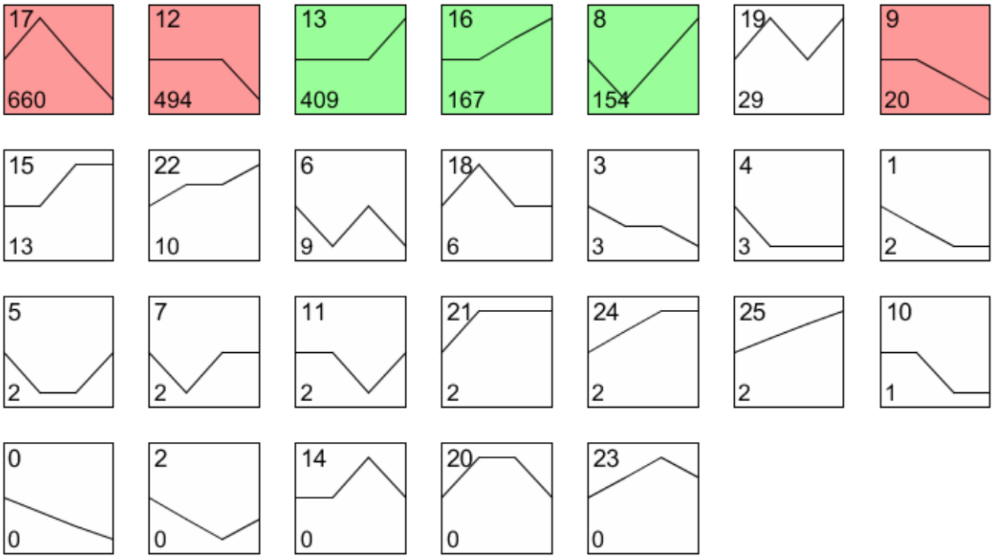
17
660
12
494
13
409
16
167
8
154
19
29
9
20
15
13
22
10
6
9
18
6
3
3
4
3
1
2
5
2
7
2
11
2
21
2
24
2
25
2
10
1
0
0
2
0
14
0
20
0
23
0

**Figure S2**
